# Vestigial-dependent recruitment contributes to robust patterning but is not required for wing-fate induction in Drosophila

**DOI:** 10.1101/2022.09.18.508456

**Authors:** Marycruz Flores-Flores, Luis Manuel Muñoz-Nava, Jeremiah Zartman, Marcos Nahmad

## Abstract

Cell recruitment is a process by which a differentiated cell induces neighboring cells to adopt its same cell fate. In Drosophila, cells expressing the wing selector gene, *vestigial* (*vg*), drive a feed-forward recruitment signal that propagates Vg expression suggesting that the Vg pattern is established as a wavefront throughout the wing pouch. However, the dynamics of Vg pattern formation does not meet this expectation. Here we show that an induction signal may drive Vg expression without Vg feedforward recruitment several cells away from the dorsal-ventral (DV) boundary. Particularly, when Vg expression is strongly inhibited either at the DV boundary or away from it, the activation of the *vg* Quadrant Enhancer still occurs at a distance, although the levels and precision of the pattern are compromised. Using live imaging of a dual-fluorescent reporter system, we show that contact-dependent cell recruitment does occur in wild-type conditions suggesting that a combination of a long-range induction signal and a signal-relay recruitment establishes Vg patterning in a robust manner. Overall, our findings reveal a previously unidentified role of cell recruitment as a robustness-conferring patterning mechanism.

**Summary Statement:** Robust formation of the Vg pattern is established by two mechanisms: a fast, long-range, but noisy induction signal; and a more precise signal relay, contact-dependent, cell recruitment process.

## Introduction

Developmental patterning is determined by cell-autonomous and non-cell-autonomous induction signals that establishes cell proliferation, differentiation, and morphogenesis (Perrimon et al., 2012). A particular case of induction is cell recruitment, in which a differentiated cell *recruits* its neighbors to differentiate into the same type as itself (Muñoz-Nava et al., 2020). Although cell recruitment is a widespread phenomenon in many developmental contexts such as the mammalian inner ear, thyroid, and kidney, and in the Drosophila wing (Kiernan, 2013; Lindström et al., 2018; Nilsson and Fagman, 2017; Zecca and Struhl, 2007a), the details about how recruitment signals regulate patterning and growth of populations of differentiated cells remains largely unknown. Moreover, the objective of cell recruitment as a developmental mechanism remains unclear. Is there a particular advantage of cell recruitment over other induction mechanisms? Particularly, why to employ cell recruitment as a patterning mechanism when, in principle, the same output may be achieved through classical morphogen patterning?

During the development of imaginal discs in *Drosophila*, wing fate is specified by the expression of the selector gene, *vestigial (vg). vg* knockout results in loss of wing structure (Williams et al., 1991; Williams et al., 1993) whereas its overexpression in other imaginal discs induces their transformation into wing-like tissue (Baena-López and García-Bellido, 2003; Kim et al., 1996). Vg patterning is a complex process that requires the integration of several signaling pathways. First, the morphogens that pattern the orthogonal axes of the wing, Wingless (Wg) and Decapentaplegic (Dpp), that emanate from the dorsal-ventral (DV) and anterior-posterior (AP) boundaries, respectively, establish a region of competence for *vg* expression and wing-fate differentiation (Kim et al., 1996; Klein and Arias, 1998; Williams et al., 1994; Zecca and Struhl, 2021). Cells at the DV boundary express *vg* through the Boundary Enhancer (*vg*^BE^) in response to Notch signaling (Irvine and Vogt, 1997; Kim et al., 1996; Williams et al., 1994). Then, the Vg pattern presumably expands through a cell recruitment signal in which Vg-expressing cells induce neighboring undifferentiated cells to express *vg* and establish the wing fate (Zecca and Struhl, 2007a; Zecca and Struhl, 2007b). The molecular identity of this cell recruitment signal is the polarization of two protocadherins, Fat (Ft) and Dachsous (Ds), which interact in a heterotypical manner at the plasma membranes of adjacent cells (Zecca and Struhl, 2010). In early wing disc development, Ft-Ds complexes are uniform and randomly distributed throughout the tissue. However, Vg initiates the recruitment signal by inhibiting *ds* transcription, thereby creating an asymmetry in Ft-Ds localization. This polarization leads to the inactivation of the Hippo pathway in the neighboring cells, thereby importing Yorkie (Yki) to the nucleus and permitting its transcriptional activity on the *vg* Quadrant Enhancer (*vg*^QE^; Zecca and Struhl, 2010; Goulev et al., 2008). This feedforward system is expected to self-propagate until no competent cells are available for recruitment.

The Vg-dependent nature of the cell recruitment signal suggests that the Vg pattern should propagate as a wavefront. However, prior work examining the dynamics of Vg patterning show a smooth gradient of Vg expression (Baena-López and García-Bellido, 2003; Muñoz-Nava et al., 2020) which is not consistent with a wavefront mode of propagation. Here, we investigate whether the formation of the Vg gradient and induction into the wing fate indeed require a Vg feed-forward input. First, we reduced Vg expression at the DV boundary and found that it affects cell survival, but when these cells were rescued from cell death, a Vg-independent signal is capable of driving Vg expression in a nearly normal pattern, although the wing pouch and the resulting adult wings are significantly diminished (Fig. 1). Furthermore, when Vg expression is knocked down in the whole dorsal compartment, *vg*^QE^ expression still propagates nearly as far as in wild-type discs, although at reduced levels (Fig. 2). This reveals that a signal that activates the *vg*^QE^ may induce Vg expression at a distance independently of feed-forward cell recruitment (Fig. 3). Although these results suggest that Vg is not absolutely required for long-range Vg expression, live imaging reveals that a contact-dependent cell recruitment process does occur in wild-type conditions (Fig. 4). Taken together, we propose that Vg patterning is established by two signaling mechanisms: a long-range induction signal that likely provides Vg patterning its graded nature; and a short-range, Vg-dependent recruitment signal that ensures a finetuning of the Vg pattern.

**Figure 1.**
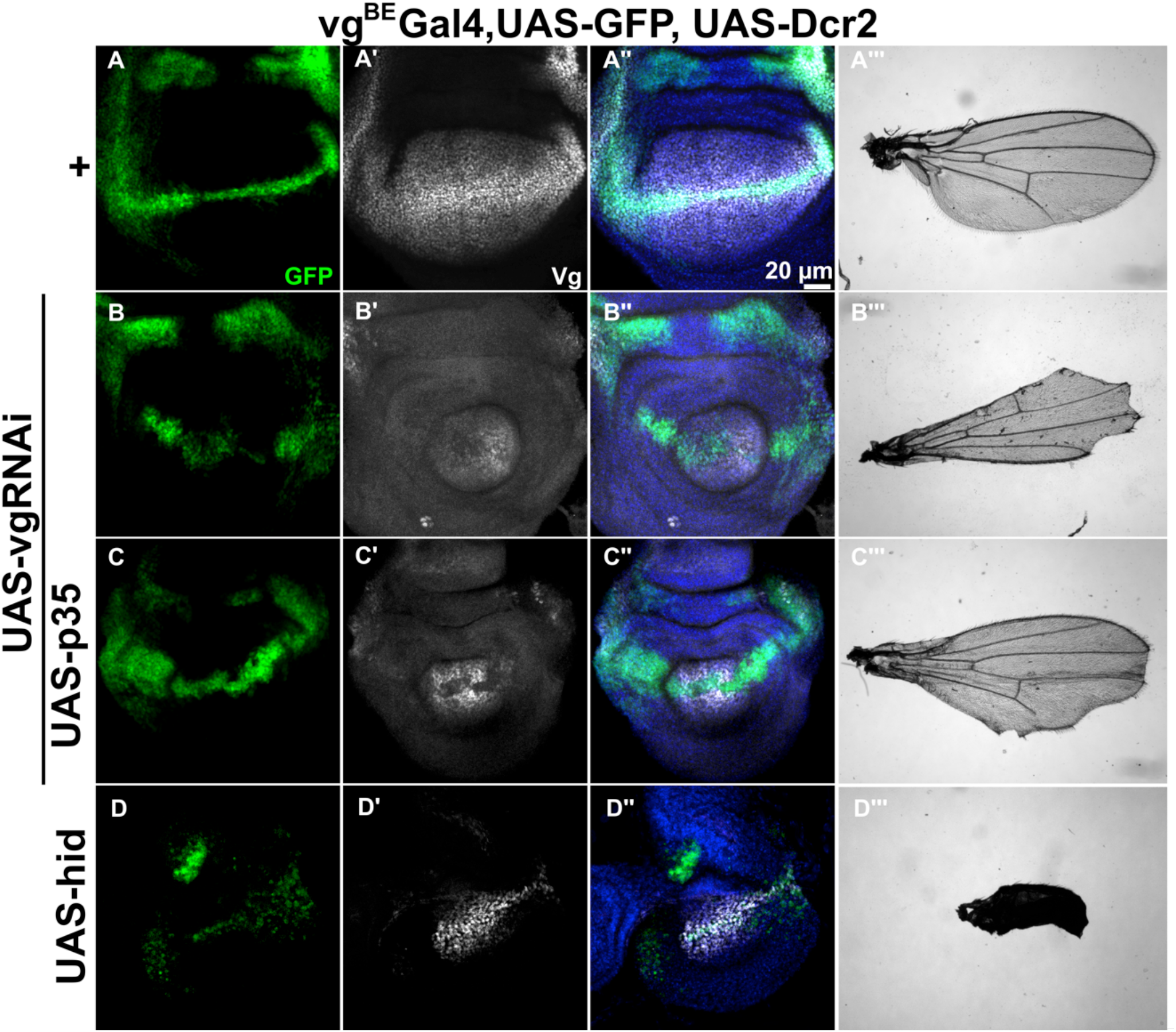
Inhibition of Vg at the DV boundary affect cell survival and growth of the wing pouch and adult wing, but is not required for long-range Vg induction. (A-D, A’-D’, A’’-D’’) Representative images of third-instar wing imaginal discs stained with GFP (green), Vg (white, using a Vg antibody), and DAPI (red) that ectopically express at the DV boundary (*vg*^BE^ domain using the Gal4-UAS system): GFP and Dcr2 (A-A’’); GFP, Dcr2, and *vg*RNAi (B-B’’); GFP, Dcr2, *vg*RNAi, and p35 (C-C’’); or hid (D-D’’). In all images, anterior is to the left and dorsal is up. (A’’’, B’’’, C’’’, D’’’) Representative adult wings that result from the corresponding genotypes.

**Figure 2.**
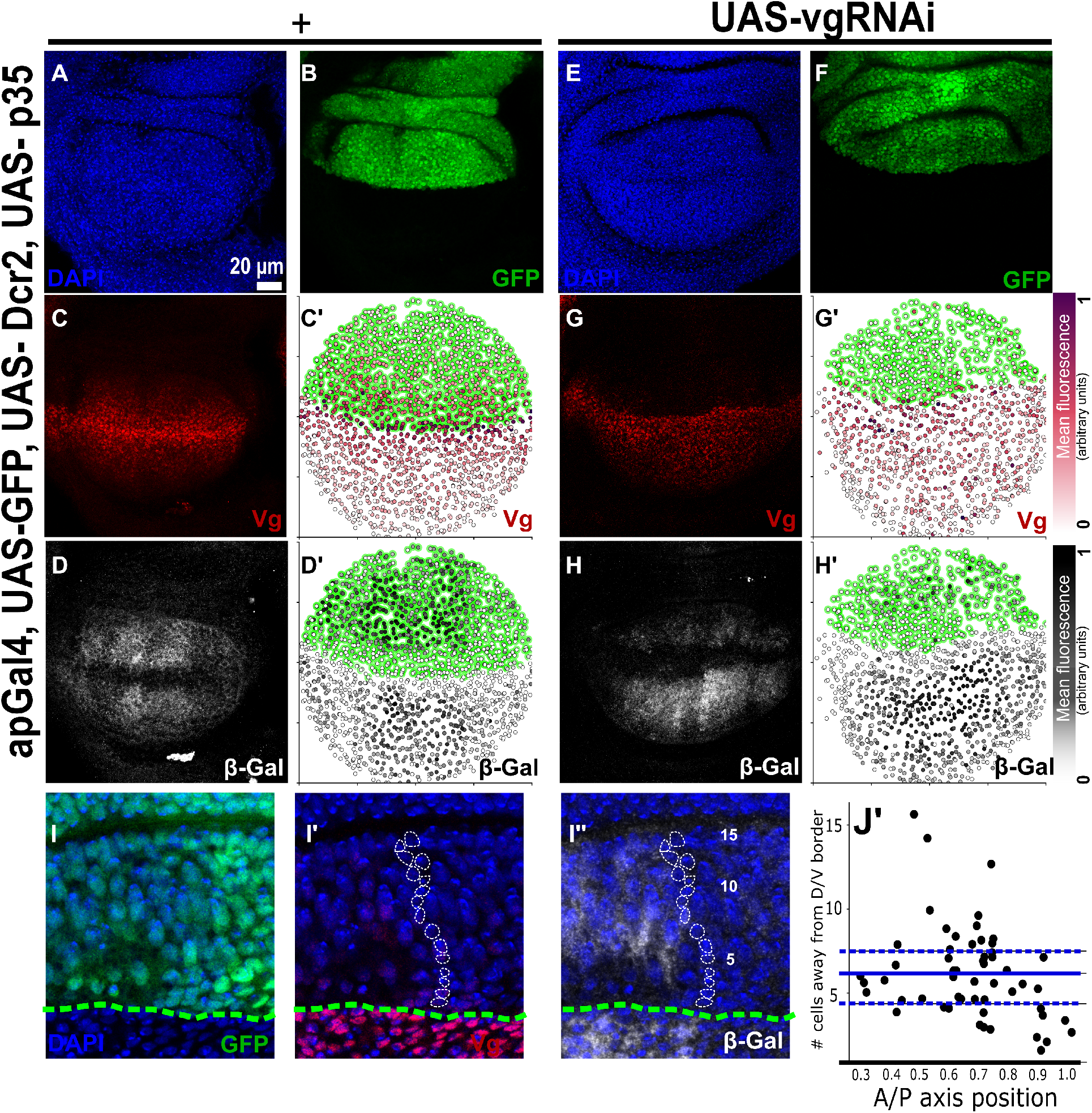
Cell recruitment signal can propagate throughout the wing pouch without a Vg-dependent feed-forward mechanism. (A-H’) Representative images of third instar wing imaginal discs stained with DAPI (A, E), GFP (B, F), Vestigial (C, G), and β-Gal (D,H) antibodies, either in a wild-type (A-D) or a Vg-knockdown (E-H) background in the dorsal compartment. (C’, G’, D’, H’) Qualitative representation of nuclear Vg or cytoplasmic β-Gal fluorescence signal, respectively. (I-I”) Representative enlargement view depicting a representative disc of the same genotype as (E-H) showing a path of cells from the DV border (marked with DAPI expression in blue) to illustrate the range of the recruitment signal (I’’, delineated in white). The DV border (bottom green-dotted line) was determined using GFP expression (I). (J’) Quantification of number of cells from the DV border that express at least half-maximal levels of β-Gal. Blue line shows the mean; Q1 and Q3 are represented by blue dotted lines). Control discs (n=9); *vg*RNAi knockdown discs (n=6).

**Figure 3.**
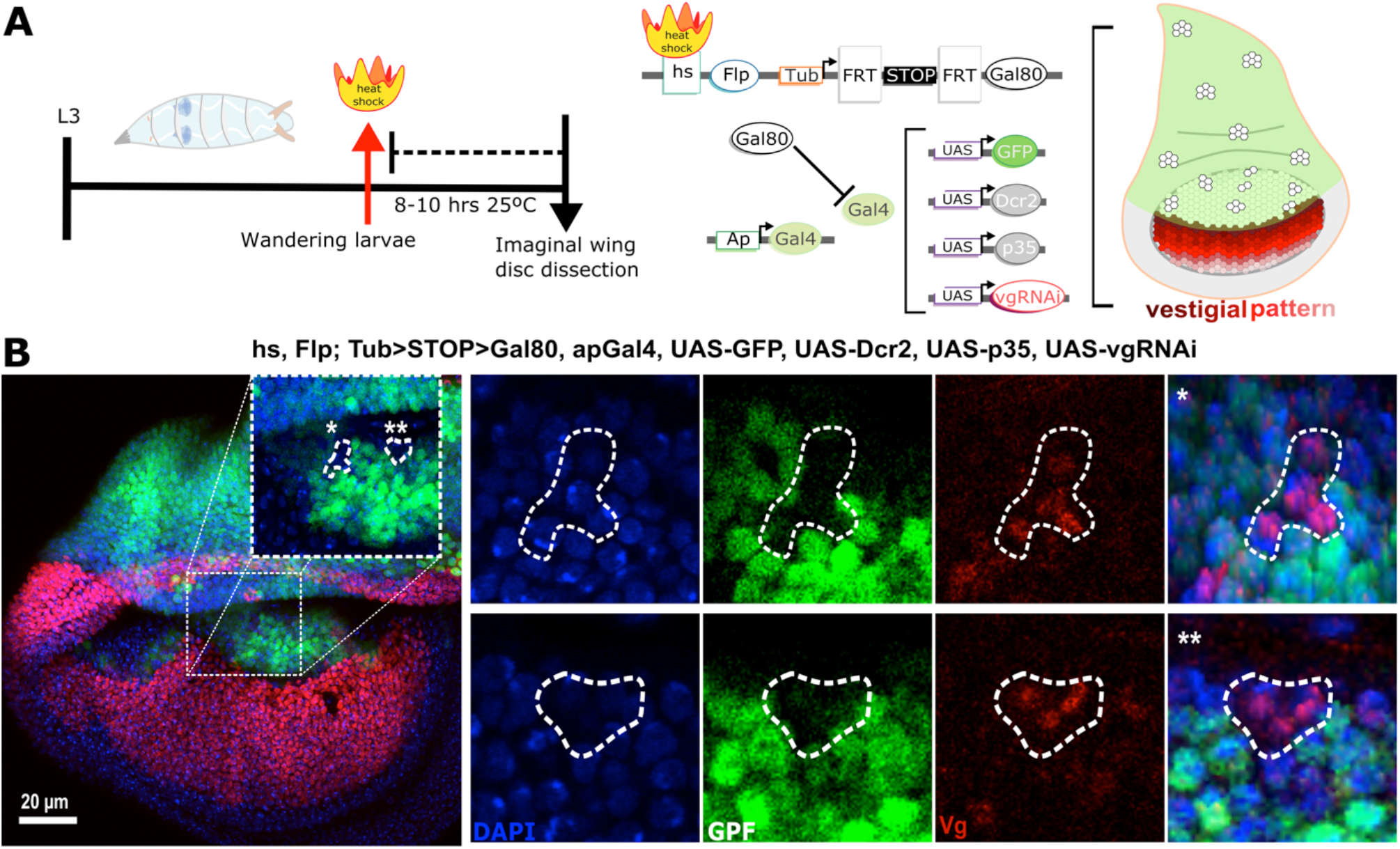
A Vg-independent induction signal activates Vg expression at a distance. (A) Experimental design for the generation of Gal80-positive clones using a heat-shock FLP-mediated recombination in *ap*-Gal4-UAS-GFP, p35, Dcr2, *vg*RNAi wing discs (Vg pattern, red). Mid-third instar larvae were heat-shocked for 10 minutes at 38°C (red arrow) and fixed/dissected 8-10 h after (late third instar, black arrow). Before heat-shock treatment, an FRT-STOP-FRT cassette prevents Gal80 to be expressed ubiquitously (under the regulation of a *tubulin* promoter). However, after heat shock, the FLP recombinase will be able to remove the FRT-STOP-FRT cassette in some cells, creating mosaics of Gal80 expressing cells. If a mosaic is located in the dorsal compartment (white clones in the disc cartoon), it prevents Gal4 from activating expression of GFP, p35, Dcr2, and *vg*RNAi (p35 and Dcr2 are expressed to prevent cell death and potentiate the RNAi effect, respectively). (C) Images of wing discs of the experimental design shown in A, stained with a Vg antibody (red), GPF (green) and DAPI (blue). Insets show the magnification of an area depicting two mosaics at different locations from the DV boundary.

**Figure 4.**
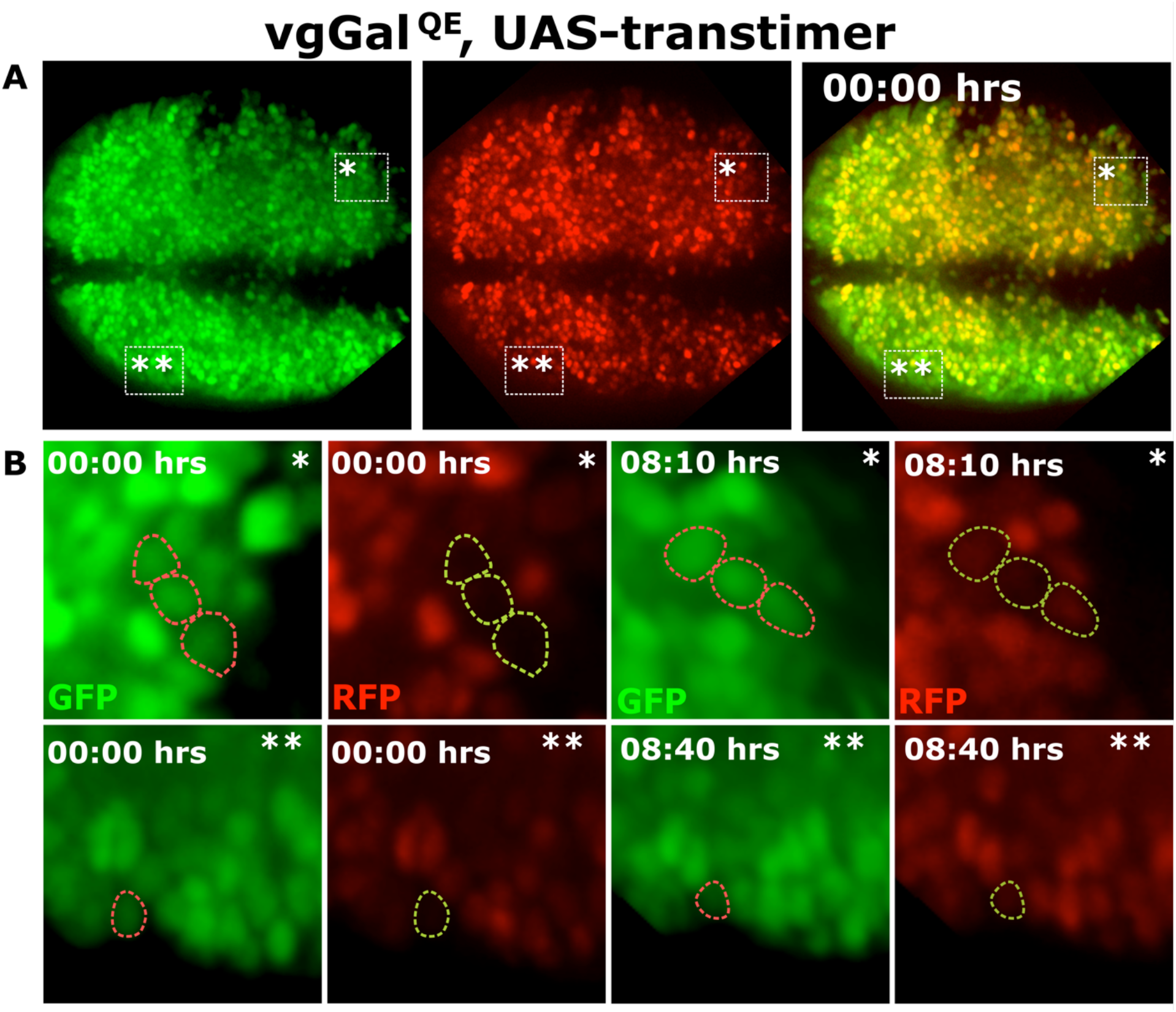
Live imaging reveals a contact-dependent cell recruitment process-taking place in the wing pouch. (A) Third instar ex vivo growing wing discs that express the Transtimer reporter under *vg*^QE^-Gal4 control. Wing disc was recorded for ~10 h using a microfluidic device with perfusion of organ culture media. Asterisks show specific places where cell recruitment is observed. (B) Two different enlarged regions from A. GFP and RFP channels at 0 h, 8:10 h or 8:40 h showing red and/or green dotted circles surrounding cells.

## Results and Discussion

### Vg expression at the DV boundary is required for survival of inducer cells, but is not essential for long-range induction of Vg

Vg is expressed at the DV boundary through the *vg*^BE^ (Fig. 1A; Kim et al., 1996; Williams et al., 1994). Thus, we asked if Vg initiates recruitment from the DV boundary, *i.e*., if Vg expression at the DV boundary was necessary to propagate its own expression in the rest of the wing pouch (Fig. 1A’, A”). To test this, we expressed a *vg*RNAi (potentiated by Dicer2 [Dcr2]) under a *vg*^BE^ -Gal4 driver to down-regulate Vg expression at the DV boundary (marked with a UAS-GFP reporter). We found that this experiment not only strongly reduced Vg expression at the DV boundary, but also affected Vg expression in the rest of the wing pouch (compare Fig. 1A’ to Fig. 1B’). We also observed that the size of the wing pouch area was dramatically reduced (compare Fig. 1A” to Fig. 1B”) and much of the distal adult wing (including the wing margin) was missing (compare Fig. 1A”’ to Fig. 1B”’). While this result supports a role for Vg in recruiting and promoting growth in the rest of the tissue, we also noticed in this experiment that the GFP reporter was mostly missing from the DV boundary within the wing pouch (compare Fig. 1A to 1B). This suggests that these cells undergo apoptosis without Vg expression, in agreement with previous studies (Baena-Lopez and García-Bellido, 2006; Delanoue et al., 2004). Indeed, when survival of these cells is ensured by co-expressing the anti-apoptotic protein p35, the GFP reporter expression along the DV boundary is recovered (Fig. 1C) and high level of Vg expression is observed in the wing pouch, except at the DV boundary itself as expected from to the ectopic expression of the *vg*RNAi (Fig. 1C’). Remarkably however, despite the rescue of GFP and Vg expression within and away from the DV boundary, respectively, the size of the wing pouch and adult wing remains strongly reduced (Fig. 1C”, 1C”’). This result confirms that Vg expression at the DV boundary is needed for cell survival and tissue growth (Delanoue et al., 2004; Pérez et al., 2011), but not for Vg induction away from the DV boundary. In fact, this interpretation is further supported by an experiment in which cells at the DV boundary are eliminated by the overexpression of *hid* that triggers apoptosis (Fig. 1D). In this case, the pattern of Vg in the wing pouch is dramatically affected (Fig. 1D’) and the resulting adult wing resembles a *vg* mutant (Williams et al., 1993; Fig. 1D”’). Taken together, we conclude that cells at the DV boundary depend on Vg to survive and proliferate, but not for Vg induction in the rest of the pouch, arguing against the hypothesis that Vg patterning is initiated by cell recruitment with a source of Vg at the DV boundary.

### Vg knockdown in the dorsal compartment does not prevent the activation of the *rg*^QE^

In the previous experiment, we cannot rule out that some Vg-expressing cells remain because the *vg*^BE^-Gal4 driver is not be strong enough to affect all the cells at the DV boundary (see overlap between GFP and Vg in Fig. 1C”). This raises the possibility that cells abutting the DV boundary propagate Vg expression throughout the disc as a feed-forward relay mechanism, *i.e*., a Vg-expressing cell recruits its neighbors to express Vg and propagate the pattern in a contact-dependent manner. Prior work suggests that the Vg-dependent feed-forward signal works through the activation of the *vg*^QE^ (Zecca and Struhl, 2007a; Zecca and Struhl, 2010). Thus, we investigated if propagation of a *vg*^QE^lacZ reporter depends on Vg expression. With this aim, we expressed a *vg*RNAi under *apterous* (*ap*)-Gal4 control, which knocks Vg expression down in all cells of the dorsal compartment (marked with a GFP reporter, Fig. 2A-D). To potentiate the effect of the RNAi and to avoid cell death resulting from the lack of Vg, we also coexpressed Dcr2 and p35, respectively. As expected, Vg is nearly eliminated in all cells of the dorsal compartment (compare Fig. 2G, G’ to Fig. C, C’). In contrast, the *vg*^QE^lacZ reporter was detected in a broad domain in the dorsal compartment (Fig. 2H, H’), reaching up to 15 cells away from the DV boundary (Fig. 2I-I”, J). However, β-Gal expression levels resulting from the *vg*^QE^lacZ reporter in Vg-knockdown discs were significantly lower and did not pattern the dorsal region of the wing pouch to the same extent as in control discs (compare Fig. 2D to Fig. 2H). This experiment strongly suggests that, at least partially, *vg* patterning may be driven by an induction signal independently of a Vg feed-forward recruitment signal. Since β-Gal expression levels were lower than normal in Vg-knockdown discs, we also considered the possibility that β-Gal expression resulted from Vg levels that remained after the Vg knockdown. To evaluate this possibility, we quantified nuclear Vg and cytoplasmic β-Gal expression (Fig. S1), and compared the distributions of the Vg to β-Gal ratios in the dorsal compartment of Vg-knockdown and control discs. We found that **residual levels of Vg remaining from *vg*RNAi expression do not explain** the distribution of β-Gal expression (Fig. S2).

The signal that drives the *vg*^QE^ in this experiment is likely emanating from the DV boundary because when an *engrailed (en)-Gal4*, which drives expression in the posterior compartment, is used instead of *ap-Gal4*, no *vg*^QE^lacZ was detected in the whole posterior compartment (Fig. S3). Since the DV boundary is a source of Wg expression (Couso et al., 1994; Neumann and Cohen, 1996) and Wg overexpression experiments expands the Vg pattern (Parker et al. 2020), Wg signaling emanating from the DV boundary is the most likely candidate to drive expression of *vg*^QE^lacZ in the absence of a Vg-dependent feed-forward signal (Fig. 2H). However, a previous study identified that in wild-type conditions, the range of Wg signaling does not exceed 11 cells (Chaudhary et al., 2019), suggesting that in addition to Wg signaling other induction signals contribute to Vg patterning. For instance, computational studies suggest that polarization of Ft-Ds signaling initiated at the DV boundary may propagate even in the absence of a Vg feed-forward reinforcement (Wortman et al., 2017).

Our results refine the model by Zecca and Struhl in which the Wg and Dpp morphogens controls Vg patterning by fueling a Vg-dependent feed-forward mechanism through the *vg*^QE^ (Zecca and Struhl, 2007b; Zecca and Struhl, 2021). In particular, we propose that Wg (and possibly Dpp, although we do not see evidence of *vg*^QE^lacZ activation when Vg is knocked down in the posterior compartment, Fig. S3) together with a Ft-Ds polarization signal act non-autonomously to drive *vg*^QE^lacZ expression in the absence of a Vg-dependent feed-forward recruitment, but not to the extent of normal Vg patterning. Nonetheless, the graded nature of the Vg pattern (rather than simply a moving wavefront) likely depends on this Vg-independent induction.

### The Vg-independent induction signal is sufficient to activate Vg expression

Our previous experiment shows that *vg*^QE^lacZ expression can be detected several cells away from the DV boundary in a Vg-independent way, but the signal response is reduced compared to a wild-type control (Fig. 2). Therefore, it remains unclear whether this Vg-independent signal is sufficient to activate Vg expression strong enough to induce them into the wing fate. To test this, we used the FLP-FRT system to generate mosaics expressing Gal80 (which inactivate the Gal4 system) surrounded by cells that express *vg*RNAi, p35, and Dcr2 under the control of *ap-Gal4* (Fig. 3A). As in our prior experiment, cells in the dorsal compartment are unable to propagate a contact-dependent feed-forward recruitment signal, but may express Vg in Gal80-expressing mosaics if the induction signal is strong enough to do so. Indeed, Vg is expressed at high levels in Gal80-expressing clones (Fig. 3B) even when they are isolated and located several cells away from the DV boundary (clones are marked by a loss of GFP; see mosaic 2 in Fig. 3B). Therefore, cells may turn on Vg expression away from the DV boundary, even when other Vg-expressing cells do not surround them. This result argues against the hypothesis that Vg induction only takes place in a contact-dependent manner and challenges the idea that Vg expression propagates as a relay process in which cells become recruiters only when they receive enough Vg to acquire the wing fate themselves (*i.e*., the threshold of Vg levels to initiate the propagation of the recruitment signal should be at least as the threshold needed to differentiate into the wing fate). How can we be sure that *vg*RNAi-expressing cells in our experiments are not sufficient to drive wing fate? Are the low, residual levels of Vg that remain from *vg*RNAi repression sufficient to drive Vg-mediated wing-fate differentiation? We argue against this case since in our *en*-Gal4, UAS-*vg*RNAi experiment the posterior portion of the adult wing is missing (Fig. S2). Taken together, we conclude that Vg patterning may be established in the absence of a contact-dependent Vg feed-forward signal.

### Live imaging reveals active recruitment at the edges of the Vg pattern in wild-type conditions

Our prior results suggest that the Vg could be driven by a Vg-independent induction signal, but does not rule out that a Vg-dependent, contact-dependent recruitment mechanism still contributes to the Vg pattern in wild-type conditions. To test this, we used live imaging of a late third-instar wing disc cultured *ex vivo* that expresses a dual-color fluorescent reporter, known as Transtimer; which produces a rapid, unstable GFP and a slow, stable RFP reporter (He et al., 2019). By driving the Transtimer reporter under *vg*^QE^Gal4 control, we were able to distinguish *yellow* (RFP+GFP) cells that established *vg*^QE^ expression several hours ago from *green* (GFP only) cells that were newly recruited (Fig. 4A). Indeed, while most cells were expressing both reporters (*yellow*), there were *green* cells at the edges of the Vg pattern (* and ** in Fig. 4). This result was consistent with a previous result in fixed tissue (Muñoz-Nava et al., 2020), but provides additional insights. First, this experiment confirms that *green* cells eventually turned into *yellow* cells, providing evidence that these cells were effectively recruited (Fig. 4B). Second, we did not see evidence of groups of *green*-only cells (*i.e*., that are not *yellow*), surrounded by other *green*-only cells, supporting the idea that this Vg-dependent cell recruitment process is a contact-dependent signal.

Our results confirm that cell recruitment does operate *in vivo* in this system and suggest that it establishes the Vg pattern together with a Vg-independent induction signal. The presence of this induction signal likely explains the graded nature of the pattern, while the recruitment signal ensures a complete pattern throughout the wing pouch. Why does pattern formation in this system uses both mechanisms, when any of them could support Vg patterning on their own? The induction signal produces a rapid, yet not very robust pattern, while a signal relay recruitment process offers a precise, yet slow patterning mechanism. By combining both, the induction signal initiates patterning by ‘reserving’ a population of cells into the wing fate while maintaining them alive and actively proliferating, whereas the recruitment mechanism ensures that all cells attain sufficient levels of Vg expression to ‘complete’ the wing specification domain. Together, these signals offer a layered mechanism that ensures robust specification of cell fates, extending the repertoire of systems in which feed-forward loop regulatory networks provide robustness and canalization in developmental systems (Ducuing et al., 2015; Le and Kwon, 2013).

## Materials and Methods

### Drosophila stocks and crosses

Unless indicated otherwise, all flies and crosses were reared on standard culture medium at 25°C.

The following stocks were used:

Fig. 1 (stocks): *vg^BE^*-Gal4 (on II; Bloomington Drosophila Stock Center [BDSC] #8222); UAS-GPFn (on III; BDSC #4776); UAS-*vg*RNAi (on II; Vienna Drosophila Resource Center # 16896); UAS-p35 (on I; BDSC #6298); UAS-Dcr2 (on III; BDSC #90938); UAS-hid (on II; BDSC #65408).

Fig. 1 (crosses): *vg^BE^*Gal4; UAS-GPFn/SM5; TM6B, Tb × UAS-Dcr2 (Control, first row); *vg^BE^*Gal4; UAS-GPFn/SM5; TM6B, Tb × UAS-*vg*RNAi; UAS-Dcr2/SM5; TM6B, Tb (second row); UAS-p35; *vg^BE^*Gal4; UAS-GPFn/SM5; TM6B, Tb (female) × UAS-*vg*RNAi; UAS-Dcr2/SM5; TM6B, Tb (third row); *vg^BE^*Gal4; UAS-GPFn/SM5; TM6B, Tb × UAS-hid (fourth row).

Fig. 2 (stocks): *ap*-Gal4 (on II; BDSC #56807); UAS-GPFn (on II; BDSC #4775)/SM5; *vg*^QE^lacZ (on III; a gift from G. Struhl, Columbia U., USA); UAS-*vg*RNAi (same as in Fig. 1); UAS-Dcr2 (same as in Fig. 1).

Fig. 2 (crosses): UAS-p35; *ap*-Gal4, UAS-GFPn; UAS-Dcr2/SM5; TM6B, Tb (female) × Sco; *vg*^QE^lacZ/SM5; TM6B, Tb (control, first and second columns); UAS-p35; *ap*-Gal4, UAS-GFPn; UAS-Dcr2/SM5; TM6B, Tb (female) × UAS-*vg*RNAi; *vg*^QE^lacZ/TM6B, Tb (control, third and fourth columns).

Fig. 3 (stocks): hs-FLP (on I; BDSC #1929); *ap*-Gal4 (same as in Fig. 2); UAS-GPFn (same as in Fig. 2); Tub-FRT-STOP-FRT-Gal80 (on III; BDSC #5145); UAS-p35 (same as in Fig. 1); UAS-*vg*RNAi (same as in Fig. 1); UAS-Dcr2 (same as in Fig. 1).

Fig. 3 (cross): *y*, *w*, *hs*-FLP; *ap*-Gal4, UAS-GPFn; UAS-Dcr2/SM5; TM6B, Tb × UAS-p35; UAS-*vg*RNAi (same as in Fig. 1); Tub-FRT-STOP-FRT-Gal80/ SM5; TM6B, Tb (select female larvae only; see Fig. 3A).

Fig. 4 (stocks): *vg*^QE^Gal4 (on III; BDSC #8229). UAS-Transtimer (on II; a gift from L. He and N. Perrimon, Harvard U., USA); TubGal80^ts^ (on III; BDSC #7017).

Fig. 4 (cross): *vg*^QE^-Gal4 × UAS-Transtimer; TubGal80^ts^/SM5; TM6B, Tb.

Fig. S3 (stocks): *en*-Gal4 (on II; BDSC #56807), UAS-GPFn (same as in Fig. 2); *vg*^QE^lacZ (same as in Fig. 2); UAS-*vg*RNAi (same as in Fig. 1); UAS-Dcr2 (same as in Fig. 1).

Fig. S3 (crosses): UAS-p35; *en*-Gal4, UAS-GFPn; UAS-Dcr2/SM5; TM6B, Tb (female) × Sco; *vg*^QE^lacZ/SM5; TM6B, Tb (control, first row); UAS-p35; *en*-Gal4, UAS-GFPn; UAS-Dcr2/SM5; TM6B, Tb (female) × UAS-*vg*RNAi; *vg*^QE^lacZ/TM6B, Tb (control, second row).

### Heat-shock induction of flip-out clones

Marked clones were generated by FLP-mediated subjecting the early-third-instar larvae to a 38°C heat-shock for 10 minutes (Fig. 3A). Larvae were then shifted to 25 °C and were dissected 8-10 h later for the immunostaining protocol.

### Live imaging

PETL (polyethylene terephthalate laminate)-based microfluidic devices were loaded with supplemented Grace’s medium (Levis et al., 2019). Organs were placed on the outlet and drawn into the device by negative pressure while suspended in supplemented Graces medium. Media flowing at 1 μl/h was used throughout imaging in the device. Using the UAS-Gal4 system, we expressed the Transtimer reporter downstream of the *vg*^QE^. During the third instar larvae, we dissected the wing disc and immediately loaded it into the microfluidic chamber, using Grace’s medium supplemented with Bis-Tris, Penn-Strp, and FBS (as described in Dye et al., 2017). Time-lapse confocal imaging was done using a Nikon Eclipse T*i* spinning disc confocal microscope (Andor). 40×/1.49-oil, and 100×/1.49-oil objectives was used for experiment. Snapshoots were taken each 10 minutes. Images were captured using MetaMorph software and image processing was performed using ImageJ and imported into R for quantification.

### Imnunostaining and confocal microscopy

Wing imaginal discs were dissected from third-instar larvae of both sexes, unless indicated. For larva dissected at a specific age AEL. After dissection in a stereoscopic microscope (Nikon SMZ800), discs were fixed in PEM-T (PEM with 0.1% of Triton X-100) with 4% paraformaldehyde, washed 3 times and blocked in PEM-T with 0.5% of BSA (Bovine Serum Albumin) for 2 h at room temperature. Then, samples were stained with primary antibodies at 4°C overnight at the following dilutions: rabbit anti-Vg (a gift from Sean Carroll and Kristen Guss, 1:200), guinea pig anti-Vg (a gift from Gary Struhl, 1:200), mouse anti-β-Galactosidase (β-Gal; 1:100). DAPI (1:1000) was used to stain nuclei. Primary antibodies were detected with Alexa Fluor 555 anti-mouse and Alexa Fluor 647 anti-guinea pig/anti-rabbit (1:1000). Imaging was done with a confocal microscope (Leica TCS SP8 Confocal Microscope) using a 63X oil-immersion objective.

### Image analysis and quantification

Wing imaginal discs were imaged as stated above. We selected three representative confocal z-stack slices in each disc; z-slices are chosen far enough so that nuclei are not double-counted. Using ImageJ, we used the DAPI channel to set a threshold and create a binarized representation. We then segment the binarized image by applying a watershed filter to obtain a nuclear segmentation mask. The image was exported into Matlab, where using the Particle Analysis function to remove particles (< 6.8 pixels). To analyze the cytoplasmic β-Gal signal, we used the final nuclear segmentation mask data and calculated the distance between all the centroids to find the minimum distance. The *Regionprops* function in Matlab was used to calculate the mean fluorescence intensity value of each nuclei or cytoplasm.

## Author contributions

Conceptualization: M.N., M.F.F.; Methodology: M.F.F.; Formal analysis: M.F.F.; Writing - original draft: M.F.F., M.N; Writing - review & editing: M.N, M.F.F., J.Z., L.M.M.N.; Supervision: M.N., J.Z.; Funding acquisition: M.N., J.Z.

## Funding

This work was supported by the Consejo Nacional de Ciencia y Tecnología of Mexico (Conacyt), [grant number CB-2014-01-236685] to M. Nahmad. L. M. Muñoz-Nava and M. Flores-Flores received financial support from Conacyt’s graduate scholarship program. ND International Mexico Faculty Grant and NIH Grant R35GM124935.

## Declaration of competing interest

No competing interests declared.

## Acknowledgments

We thank Gary Struhl for kindly providing us the guinea pig-Vestigial antibody. We also thank Jose Luis Fernandez for technical assistance, and members of the Nahmad and Zartman laboratories for interesting discussions.

## Notes

### Competing Interest Statement

The authors have declared no competing interest.

